# Glycolytic Specialization Shapes Neuronal Physiology and Function *in vivo*

**DOI:** 10.64898/2026.02.17.706437

**Authors:** Aaron D. Wolfe, Longgang Niu, L. Safak Yilmaz, Snusha Ravikumar, Matthew Thomas, Albertha J. M. Walhout, Zhao-Wen Wang, Richard H. Goodman, Daniel A. Colón-Ramos

## Abstract

Neurons perform diverse functions that impose distinct energetic demands, but how energy-metabolic pathways are matched to these functions *in vivo* remains unknown. Here we show that two functionally divergent sister chemosensory neurons in *C. elegans,* ASEL and ASER, exhibit asymmetric glycolytic flux, with ASER exhibiting high and ASEL having low levels of glycolysis. Metabolic imaging, metabolic network modeling, and electrophysiology measurements reveal that ASER’s elevated glycolysis supports a hyperpolarized resting potential, low input resistance, and rapid repolarization that enable a distinct functional role compared to ASEL. Impairing glycolysis collapses these electrophysiological specializations without abolishing neuronal excitability, and selectively disrupts ASER’s calcium responses while leaving ASEL largely unaffected. These findings demonstrate that neuron-specific glycolytic programs shape core biophysical properties and are required for functional identity *in vivo*, establishing metabolism as an active determinant of neuronal physiology.

## Introduction

Neurons show great diversity in function, and their specialized computational roles arise primarily from cell-intrinsic properties such as ion channel composition, membrane conductance, and firing patterns that all define their physiological nature^1–3^. The elements that support the inter- and intracellular communications of neurons require substantial energetic input, and as a result the brain is the most energetically-demanding organ in the body^4–6^. Because of this, impairment of brain energy metabolism can dramatically affect neuronal function or physiology^7,8^. Accordingly, during increased neuronal activity, neurons adapt their metabolic outputs to meet these elevated energetic needs^9–11^. These observations underscore a tight coupling between neuronal physiology and energy metabolism, raising the question of how metabolic programs are organized across neuron types to support their distinct functions.

Specific metabolic programs are linked to cellular functions in specific tissues. For example, M1 killer macrophages use glycolysis for energy and reprogram mitochondria for ROS generation^12^; type IIB fast twitch muscle fibers use glycolysis to support physiological demands of short bursts of rapid energy production^13^; and vertebrate photoreceptors depend on elevated glycolysis to sustain an energetically-costly ion dark current that underlies sensory transduction^14^. Whether metabolic programs differ across individual neurons to support their function is less understood. Recently, we observed that the model organism *C. elegans* displays a range of stable glycolytic states that are characteristic of specific neuron types^15,16^, suggesting metabolic specialization may be an under-appreciated property of neurons. Yet how such metabolic states are established and whether they are causally linked to cell-specific physiological functions *in vivo* remain unclear.

To investigate how distinct neuronal functions associate with baseline metabolic states at single-cell resolution, we turned to the *C. elegans* ASE chemosensory neurons, ASEL and ASER. These lateralized sister cells provide sensory input into the same neuronal circuit but adopt asymmetric physiological properties and behavioral roles^17–19:^ ASEL is excited by increases (“ON” responses) in NaCl concentration, whereas ASER is excited by NaCl decreases (“OFF” responses), enabling animals to interpret various salt gradients^20,21^. These functional asymmetries are reflected in divergent calcium dynamics, synaptic communication patterns, and contributions to chemotaxis^21–24^. Their mirrored identities, common circuit context and divergent functional specializations make ASEL and ASER an ideal system for uncovering how neuron-specific metabolic profiles are set, and how they contribute to the physiological and computational properties of single neurons *in vivo*.

## Results

### Distinct baseline glycolytic states in the functionally asymmetric ASE neurons

To examine whether the asymmetric physiology of ASEL and ASER is accompanied by differences in baseline patterns or regulation of metabolism, we established a single-cell metabolic profiling platform in the two functionally distinct ASE neurons (Fig. 1A). We achieved this via three genetically encoded biosensors that report distinct nodes of glycolytic activity *in vivo* (Fig. 1B): HYlight^25^, which measures the central glycolytic intermediate fructose-1,6-bisphosphate (FBP); R-iLACCO1.2^26^, which reports lactate production as the endpoint of glycolysis; and SoNar^27^, which monitors the cytosolic NADH/NAD⁺ ratio, a metabolic cofactor tightly coupled to glycolytic flux. By visualizing animals expressing these sensors in ASEL and ASER, we characterized their resting states of glycolysis with single-cell resolution in living animals.

**Figure 1:**
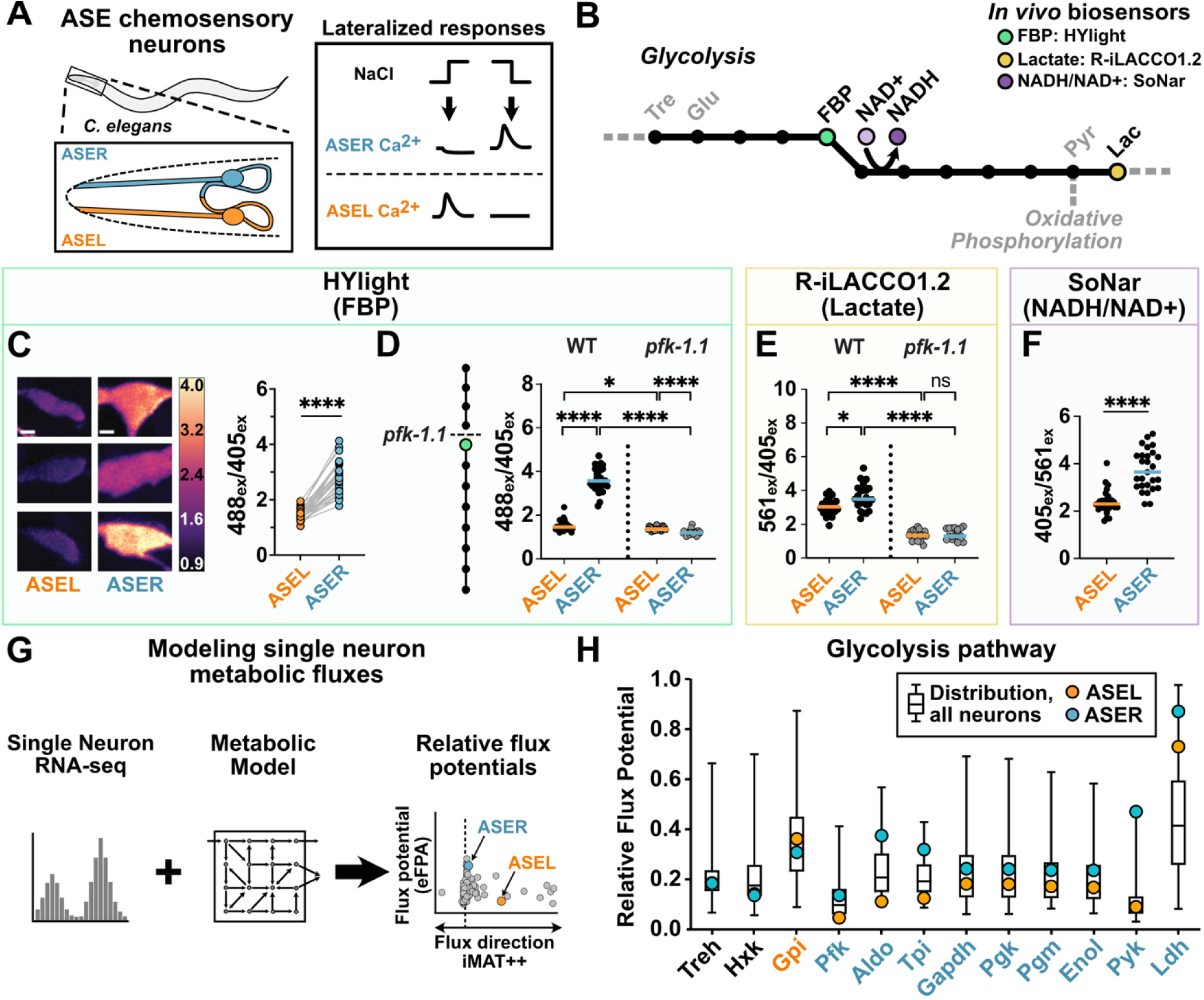
Distinct states of glycolysis in the functionally asymmetric ASE neurons. (A) Left, cartoon diagram of the location of both ASE chemosensory neurons in *C. elegans*; right, schematic of lateralized ASE calcium responses to salt changes (B) Depiction of the glycolysis pathway showing metabolites detected by the three biosensors used in this study. (C) Three representative ratiometric images of HYlight expression in ASE cell somas; ratio of excitation by 488 nm over 405 nm is displayed. Scale bar is 3 µm. Right, paired mean soma values for ASER and ASEL; each dot pair represents measurements from a single animal. (D) Mean soma measurements of HYlight in wild type vs. CRISPR-deletion mutant *pfk-1.1(ola458)*; each dot represents one animal. Horizontal line marks the median for all animals. (E) Mean soma measurements of R-iLACCO1.2 for wild-type and *pfk-1.1(ola458)* mutant animals. (F) Mean soma measurements for SoNar in ASEL and ASER in wild-type animals. (G) Diagram of the MERGE computational pipeline^32^ used to predict attainable fluxes via the iMAT++ algorithm and to estimate relative flux potentials using enhanced flux potential analysis (eFPA)^30^ in *C*. *elegans* neurons and other cell types. (H) Predicted flux potentials for reactions in the glycolysis pathway. Bar graphs represent distribution for 103 neuron classes in *C. elegans*. Blue and orange dots represent values for ASER or ASEL, respectively. Enzymes in x-axis are colored by which ASE neuron is predicted to have higher flux potential (ASER, blue; ASEL, orange; if similar, black). See table S1.\**P*<0.5; \*\*\*\**P*<0.0001; n.s., non-significant. Statistical details available in table S2.

Comparing measurements of the ratiometric FBP sensor HYLight in ASER and ASEL within the same animals revealed significantly higher HYlight signal in ASER as compared to ASEL under baseline conditions (Fig. 1C). This difference was absent when using HYlight-RA (Fig. S1A), a binding-deficient control sensor. To validate that these HYlight measurements track *bona fide* changes in glycolytic flux, we examined mutants that perturb FBP production (Fig. 1D, S1B). The genes *pfk-1.1* and *pgk-1* encode enzymes acting upstream and downstream of FBP, respectively. The allele *pfk-1.1(ola458*), a CRISPR deletion of the primary phosphofructokinase ortholog in *C. elegans*^15^, displayed a markedly reduced HYlight signal in ASER, but had minimal effect in ASEL (Fig. 1D). A loss-of-function allele of the sole phosphoglycerate kinase, *pgk-1(tm5613),* caused HYlight signal to increase in both ASEL and ASER, consistent with accumulation of upstream metabolites, including FBP (Fig. S1B). Together, these data: 1) demonstrate that HYlight accurately reports changes in glycolytic pathway activity in the ASE neurons; 2) provide a framework for interpreting the elevated FBP observed in ASER relative to ASEL; and 3) suggest that the glycolytic pathway is highly active in ASER compared to ASEL under basal conditions.

To further examine if the elevated HYlight signal in ASER reflects increased glycolytic flux, we used the reporter R-iLACCO1.2 to measure lactate, a terminal endpoint of glucose metabolism via glycolysis. We observed that ASER exhibited significantly higher lactate levels than ASEL (Fig. 1E), consistent with enhanced glycolytic output. As with FBP, we used genetic perturbations to validate the reporter’s specificity *in vivo*: *pfk-1.1(ola458)* markedly reduced R-iLACCO1.2 signal in both neurons. A CRISPR-engineered deletion mutant of the sole lactate dehydrogenase ortholog, *ldh-1(ola514)*, also reduced the R-iLACCO1.2 signal (Fig. S1C). We then measured the cytosolic NADH/NAD⁺ ratio by using the sensor SoNar, and observed a significantly higher SoNar signal in ASER as compared to ASEL (Fig. 1F), consistent with elevated glycolysis. Together, measurements with these three different biosensors for glycolysis indicate that ASER exhibits higher utilization of glycolysis compared to ASEL.

To determine whether these metabolic differences are reflected in their transcriptional expression patterns of these neurons, we modeled flux potentials. Changes in metabolic gene expression are predictive of metabolic fluxes^28–30^, so we integrated single-cell RNA-seq data for 103 neuron classes^31^ with the *C. elegans* iCEL1314 metabolic network using the MERGE computational pipeline^32^ to estimate pathway-specific flux potentials (Fig. 1G). This flux modeling predicts higher potential in ASER for all glycolytic reactions from phosphofructokinase (Pfk) to pyruvate kinase (Pyk), and lactate dehydrogenase (Ldh) as well as other glycolysis-adjacent pathways (Fig. 1H, S1D-F). The model also predicts elevated flux potentials for glycogen degradation, an observation consistent with our past report on the role of glycogen supporting metabolic plasticity in ASER^16^. Together, these modeled flux potentials support our biosensor measurements in defining ASER as a high-glycolysis neuron.

### Intrinsic neuronal identity specifies distinct glycolytic states and responses

A canonical approach for assessing differences in glycolytic pathway activity across cell types is the glycolysis stress test^33^, a metabolic phenotyping assay that reports cellular dependence on and flexibility of glycolytic energy production. To further examine how ASEL and ASER make differential use of glycolysis, we adapted key features of this *in culture* assay to an *in vivo* context by leveraging single-cell biosensors, applying transient hypoxic conditions using a microfluidic device^34^ (Fig. 2A) and monitoring the responses of the two ASE neurons simultaneously. We determined three parameters in the response of these neurons: 1) *basal glycolysis*, as the HYlight signal before hypoxia is applied; 2) *glycolytic capacity* as the HYlight signal achieved upon hypoxia; and 3) *glycolytic reserve* as the difference between glycolytic capacity and basal glycolysis. Both ASEL and ASER responded to hypoxia by increasing glycolysis, while the binding deficient HYlight-RA showed no response (Fig. 2A). Consistent with their different basal glycolytic states, we also observed that ASEL had a significantly higher glycolytic reserve compared to ASER (Fig. 2B), though ASER had a significantly higher glycolytic capacity (Fig. 2C). Together, our data indicate that ASEL and ASER both adapt glycolysis upon energetic stress, yet differ in their basal utilizations of this pathway.

**Figure 2:**
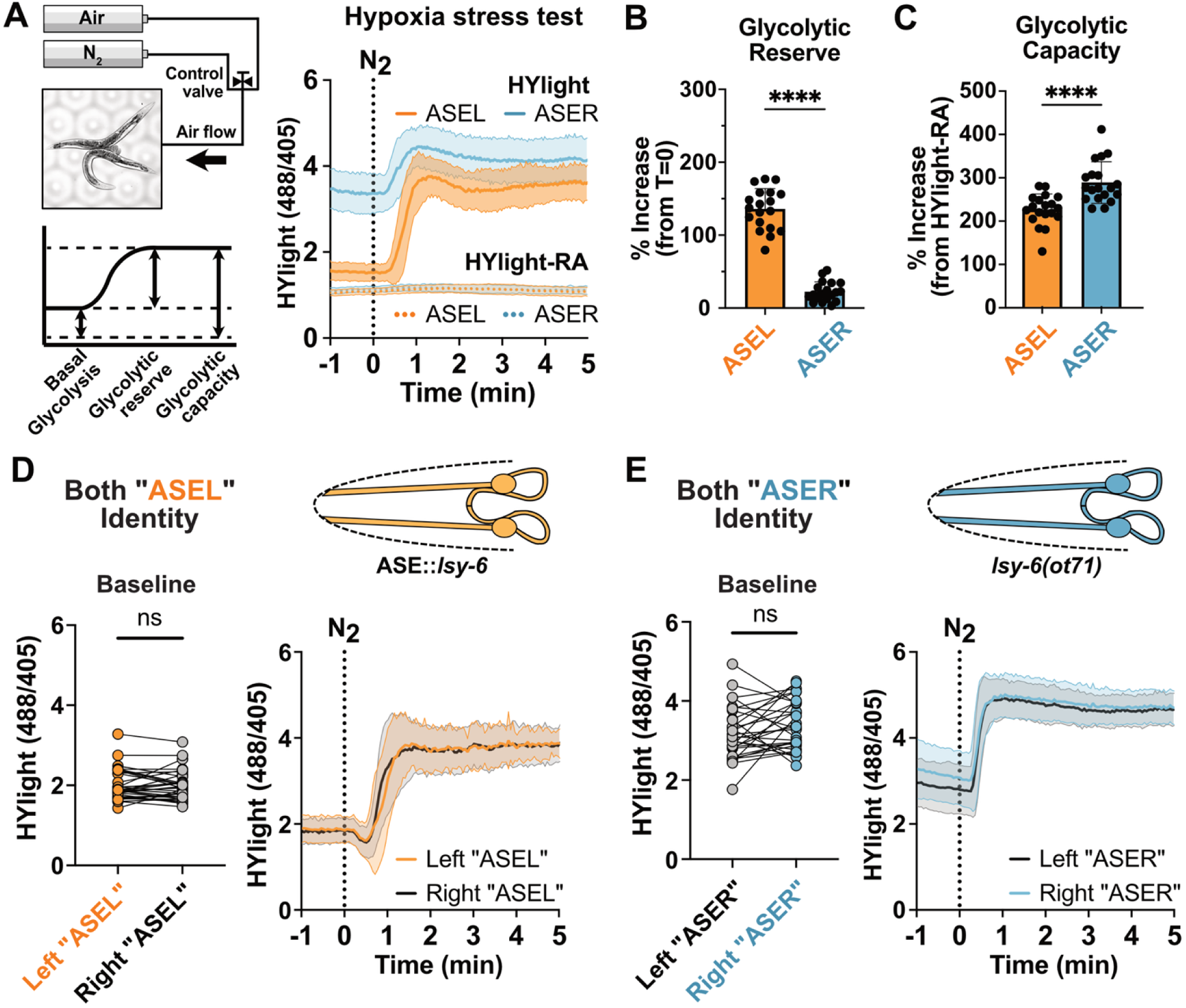
Basal usage of glycolysis is associated with neuronal identity. (A) Left, diagram of the hypoxia microfluidics system^34^ used in this experiment, with a hypoxia stress test curve and definition of key measurements shown below. Right, pooled soma measurements of HYlight (solid line) or negative control HYlight-RA (dashed line) in ASEL (orange) or ASER (blue) upon hypoxia (vertical dotted line at time 0 reflects induction of hypoxic condition). (B-C) Values for glycolytic reserve (B) and capacity (C) calculated from data in Fig. 2A. (D-E) Mean soma measurements of either the *olaIs141; otIs204* (two ASEL-identity) animals (D) or *olaIs141; lsy-6(ot71)* (two ASER-identity) animals (E). Each dot represents a paired measurement of either the left-side or right-side neuron within the same animal. Shaded region of glycolysis stress tests represents standard deviation around the mean. \*\*\*\**P*<0.0001; ns, non-significant. Statistical details available in table S2.

The ASEL and ASER cell fates are specified by a deterministic, transcriptionally encoded left-right differentiation program^17–19^. To determine whether the metabolic differences we observed are cell-intrinsic consequences of this fate-specification program, we examined mutants that disrupt the ASE transcriptional cascade (Figs. 2D-E). In *otIs204[ceh-36p::lsy-6]* transgenic animals, in which both ASE neurons adopt an ASEL-like identity^35^, the metabolic differences seen in wild-type animals were abolished. The two neurons exhibited low baseline HYlight levels akin to ASEL in wild-type animals, and both neurons displayed similar responses upon the glycolytic stress test (Fig. 2D). Conversely, in *lsy-6(ot71)* mutants, both neurons adopt ASER-like identities^36^ and exhibited HYlight baseline levels and responses that matched the ASER wild-type characteristics (Fig. 2E). Together, these results indicate that the distinct glycolytic states of ASEL and ASER arise from cell-intrinsic properties of their transcriptionally defined neuronal identities.

### Glycolysis responds to ion flux across the membrane

While glycolysis is rapidly modulated by neuronal activity^9,37–39^, we observe stable, identity-dependent differences in basal glycolytic states between ASEL and ASER. Therefore, we asked whether sensory-driven neuronal activity contributes to these metabolic differences. To do so, we built a microfluidic system^16^ that allows rapid and precise modulation of a relevant sensory stimulus for ASE neurons (salt concentrations), while performing confocal imaging of calcium, using JGCaMP8m^40^, or HYlight (Fig. 3A). We observed that ASEL and ASER exhibited robust calcium transients in response to increases (for ASEL) and decreases (for ASER) in NaCl concentration (Fig. 3B, S2A), consistent with the known functions of these cells^21^. Under the same stimulation paradigm, we observed that both neurons showed comparable changes in HYlight values upon stimulation (Figs. 3C, S2B). Our findings are consistent with prior work showing that neuronal activity can drive rapid elevation of glycolysis^9,16,37–39^. Importantly, our results reveal that the ASE neurons use two layers of metabolic modulation: a transcriptionally encoded baseline state that differs between ASEL and ASER, and a shared capacity for adaptive, activity- or stress-dependent upregulation of glycolysis, likely mediated through post-transcriptional or biochemical mechanisms.

**Figure 3:**
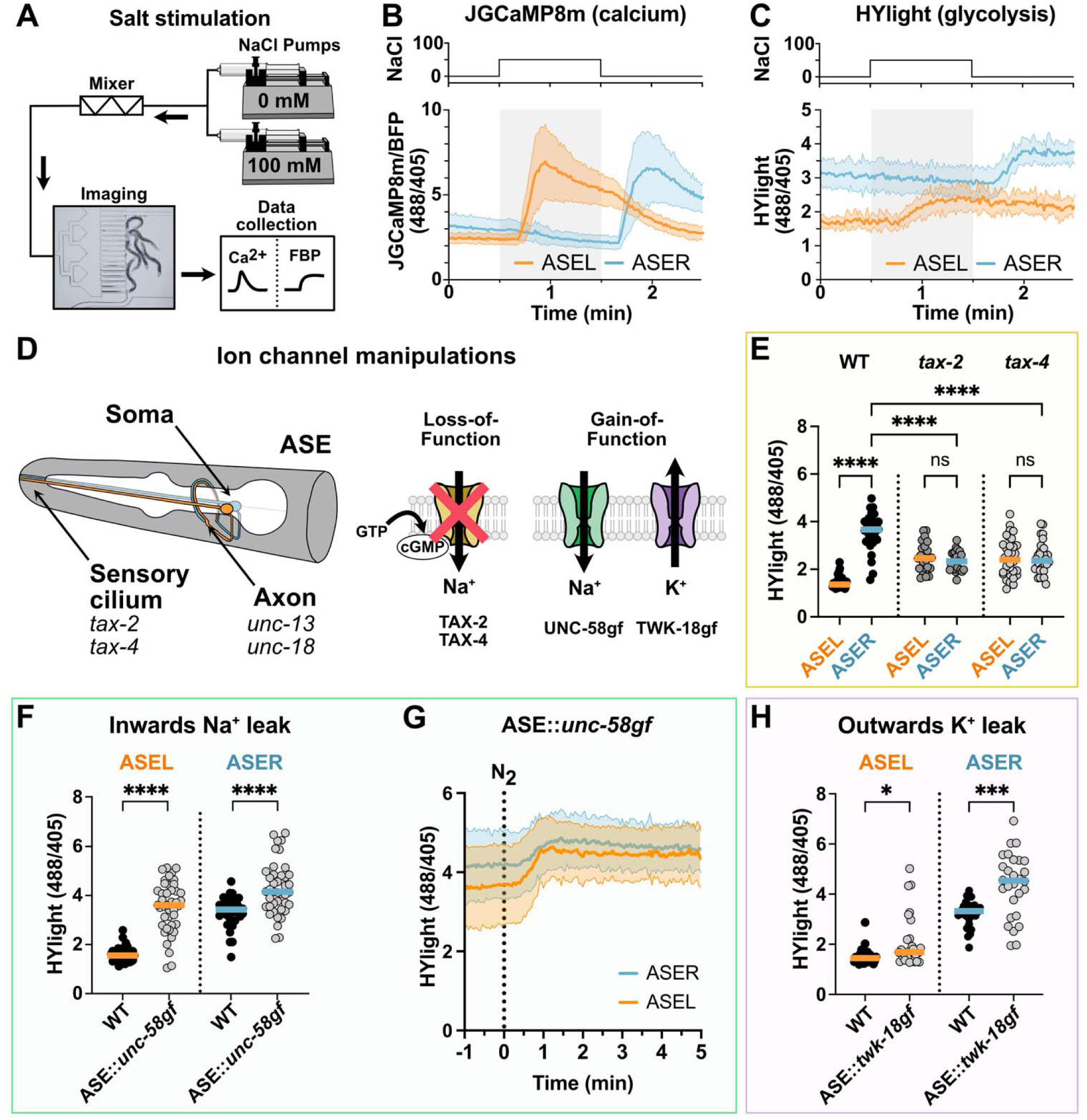
Glycolysis responds to changes in membrane ion currents. (A) Diagram of the salt mixing microfluidics system used to stimulate the ASE neurons with changing salt concentrations. (B) Pooled mean soma calcium responses (as JGCaMP8m change to BFP fluorescence) for a salt pulse experiment. Both ASEL and ASER values were captured for the same individual worms. (C) A similar experiment as in (B), but measuring HYlight responses instead. (D) Left, diagram of ASE neurons, showing the sensory cilia relative to the soma and axonal regions. The genes unc-13 and unc-18 function in the axon to facilitate synaptic vesicle release (see Fig. S2C). The genes *tax-2* and *tax-4* function in the cilia to initiate sensory responses. Right, schematic of loss-of-function experiments of the *tax-2/4* cation channels and gain-of-function experiments using heterologous expression of the hyperactive alleles *unc-58(e665)* and *twk-18(e1913)*. (E) Mean soma measurements of HYlight in either wild-type, *tax-2(gk117937),* or *tax-4(p678)* animals. (F) Mean soma measurements comparing wild-type animals and animals expressing UNC-58(L428F) in both ASE neurons. (G) Pooled HYlight measurements from a glycolysis stress test using the same animals as in (F). (H) As in (F) but expressing TWK-18(G165D) in both ASE neurons. Shaded regions around each time course reflect the standard deviation around the mean. \**P*<0.5; \*\*\**P*<0.001; \*\*\*\**P*<0.0001; ns, non-significant. Statistical details available in table S2.

We next sought to determine whether this metabolic distinction of ASEL and ASER arises from differences in basal metabolic demands. Prior studies that model the energy budget of neurons have indicated that a majority of neuronal ATP is consumed either maintaining ionic gradients for membrane potentials or supporting synaptic transmission and its associated processes^4,5,41^. Thus, we used genetic mutants to determine which of these processes might contribute to the distinct metabolic states of these neurons (Fig. 3D). We observed that loss-of-function mutations in *unc-13(s69)* and *unc-18(e234)*, which impair synaptic vesicle priming and docking respectively, had no effect on the difference in baseline glycolysis between these neurons (Fig. S2C). However, loss-of-function mutations in *tax-2* and *tax-4*, which encode the β and α subunits of the cGMP-gated cation channel required for sensory-evoked neuronal activity^21^, abolished the metabolic distinction between ASEL and ASER. (Fig. 3E). These results implicate ion channel-mediated membrane conductance, rather than synaptic transmission, as a key determinant of the basal glycolytic state of ASE neurons.

To further test whether membrane potential and ionic gradients are linked to the observed basal glycolytic states of the ASE neurons, we manipulated conductance of either sodium or potassium ions via ectopic expression of cation channels carrying hyperactive gain-of-function variants of two-pore domain channels, UNC-58 and TWK-18 (Fig. 3D, F-H). ASE-specific expression of UNC-58(L428F), which selectively increases inward sodium flux^42^, produced a significant elevation in HYlight signal in both ASE neurons (Fig. 3F). Interestingly, the differences observed for the two ASE neurons in the glycolytic stress test were abrogated in the UNC-58(L428F) animals (Fig. 3G), consistent with a role for membrane potential in setting these distinct baseline differences between ASEL and ASER. ASE-specific expression of TWK-18(G165D), which enables a constitutive outwards flow of potassium^43^, also resulted in a significant increase in the baseline of ASER, as well as a modest though significant effect in ASEL (Fig. 3H). Taken together, these experiments show that increasing the conductance of either Na+ or K+ cations across the membrane elevates the glycolytic state of ASE neurons, suggesting that differences in ion channel composition and conductance could contribute to the identity-dependent differences in basal glycolysis observed for ASEL and ASER.

### Glycolysis is necessary to maintain ASER electrophysiological and functional identity

Because differences in ion channels play an important role in setting the electrophysiological properties of neurons, we next asked whether the elevated glycolysis of ASER is derived from distinct membrane properties relative to ASEL. Using either wild-type or *pfk-1.1(ola458)* animals in which glycolysis is impaired, we performed current-clamp recordings from both ASE neurons *in situ* and recorded membrane voltage responses to incremental current injections (Fig. 4A). We observed that the resting membrane potential (RMP) was significantly more hyperpolarized in ASER compared to ASEL in wild-type animals, while this difference was lost in *pfk-1.1(ola458)* animals (Figs. 4A,B). In contrast, ASEL showed no change in RMP when comparing wild-type and *pfk-1.1(ola458)* animals. The input resistance of ASER was also significantly lower than ASEL in wild type (Fig. 4C), while this difference was also lost in *pfk-1.1(ola458)* animals. Notably, no change in membrane capacitance was observed in ASER between wild-type and *pfk-1.1(ola458)* animals (Fig. S3A), showing that the change in input resistance between them was not due to a change in cell size.

**Figure 4.**
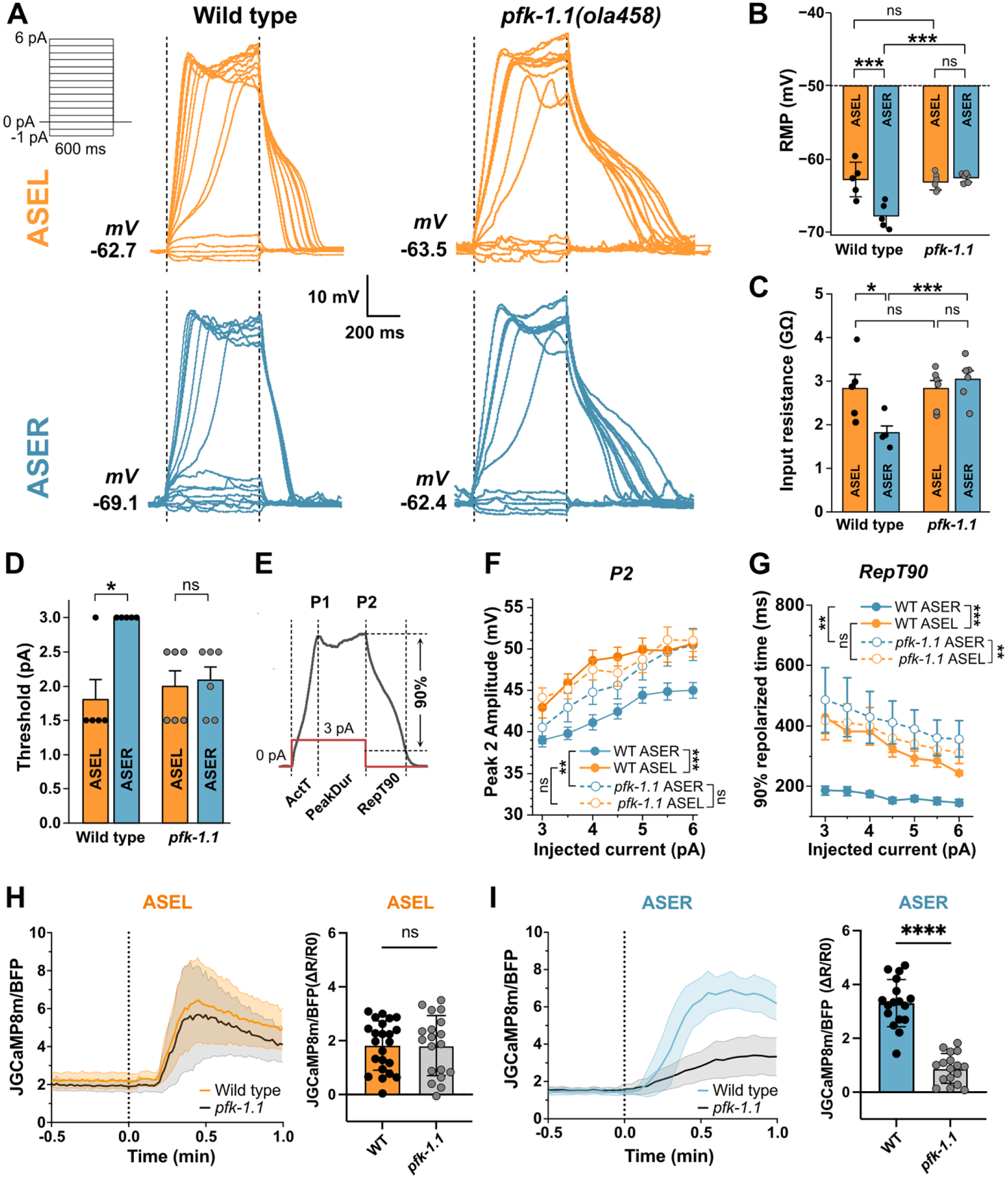
Glycolysis is necessary to support physiology and sensory responses in ASER. (A) Current injection protocol and the resulting representative traces of membrane voltage responses of ASER and ASEL in wild-type and *pfk-1.1(ola458)* animals. All subsequent analyses are shown for a sample size of five wild-type and six *pfk-1.1* animals. (B-D) Statistical comparisons of the resting membrane potential (B), input resistance (C), and threshold for active responses (D) between neurons and across genotypes. (E) A voltage response trace illustrating the definitions of Peak 1 (P1) and Peak 2 (P2) amplitudes, activation time (ActT), 90% repolarization time (RepT90), and peak duration (PeakDur). See also Fig. S3. (F-G) Comparisons of P2 amplitude (F) and 90% repolarization time between neurons and across genotypes. Data for B-G are shown as mean ± SEM. (H) Left, pooled calcium traces in ASEL upon a salt pulse (0 mM NaCl to 50 mM NaCl, at the vertical dotted line). Right, comparison of the percent increase for the calcium peak for wild-type vs *pfk-1.1(ola458)* animals relative to stimulation. Each dot represents one animal. (I) Left, pooled calcium traces observed within ASER upon a salt pulse stimulation paradigm known to activate ASER (75 mM to 25 mM NaCl). Right, the percent change as in (H, right) but for ASER data. Shaded regions around each time course reflect the standard deviation around the mean. \**P*<0.5; \*\**P*<0.01; \*\*\**P*<0.001; \*\*\*\**P*<0.0001; ns, non-significant. Statistical details available in table S2.

Consistent with its hyperpolarized RMP and lower input resistance, the ASER neuron had a significantly higher threshold for active (regenerative) responses compared to ASEL, while in the *pfk-1.1(ola458)* mutant background, this distinction was lost (Fig 4D). In addition, other physiological characteristics of the ASER responses were selectively affected in the *pfk-1.1* mutant (Figs. 4E-G, S3A-D). ASER in *pfk-1.1(ola458)* had significantly higher peak voltage amplitudes compared to ASER in wild type (Figs. 4F, S3B); this same trend was also observed for 90% repolarization time (Fig. 4G), though not observed for activation time nor peak duration (Figs. S3C-D). Notably, the threshold for regenerative responses, peak voltage amplitudes, and 90% repolarization time were unchanged in ASEL between wild-type and *pfk-1.1* animals.

Together, these findings suggest that the higher glycolytic state in wild-type ASER is necessary to maintain distinct electrophysiological properties compared to ASEL, including a more hyperpolarized RMP, lower input resistance, and higher threshold for active depolarization.

To determine whether these asymmetries in glycolysis and electrophysiology between ASEL and ASER underlie their roles in sensation, we next compared calcium responses of ASEL and ASER to stimulating changes in NaCl between wild-type and *pfk-1.1(ola458)* animals. ASEL calcium responses to NaCl were indistinguishable in wild-type and *pfk-1.1(ola458)* animals, indicating that disruption of glycolysis does not affect calcium responses in ASEL (Fig. 4H). In contrast, *pfk-1.1(ola458)* mutant worms exhibited a significantly reduced calcium response in ASER compared to wild-type worms (Fig. 4I). All together, these findings show that ASER, but not ASEL, requires glycolytic flux to support its characteristic sensory physiology, underscoring that elevated glycolysis is fundamental to the functional identity of this neuron.

## Discussion

Our findings reveal that neurons exhibit cell type-specific metabolic programs that are integral to their physiological identities. Although the two ASE neurons together act as sensory inputs to the same behavioral circuit, they have distinct roles in the computation of this behavior. Here we have shown these two cells also maintain different metabolic states that support differentiated electrophysiological properties. ASER utilizes a metabolic paradigm organized around elevated glycolysis and depends upon this pathway to support its more hyperpolarized resting membrane potential, lower input resistance, and faster membrane repolarization kinetics. Disruption of glycolysis led to a loss of these features specifically in ASER, resulting in this neuron adopting properties of the less glycolytic ASEL, and dramatically affected sensation-induced calcium responses. Our findings thus establish metabolic state as a core parameter defining neuronal identity and function, akin to other properties such as ion channel composition, neurotransmitter use, and synaptic connectivity.

Neuronal energy demands are traditionally considered in the context of activity-driven consumption, yet neurons are also predicted to incur substantial metabolic costs at rest to maintain their membrane potentials^4,5,11^. This principle is well illustrated by photoreceptors, where a dark current mediated by steady Na⁺ influx enables exquisite sensitivity but imposes one of the highest ATP burdens in the nervous system^14^. Here we have shown that increasing ion fluxes across the cell membrane by ectopic expression of hyperactive cation channels resulted in increased glycolysis, implicating passive currents in contributing to the different metabolic states. The hyperpolarized resting potential and reduced input resistance of ASER are likely indicative of greater outwards potassium flux at rest. Such elevated potassium efflux might incur higher demands on active ion pumps, such as the Na^+^/K^+^ ATPase, which is known to drive glycolysis upon neuronal stimulation^44^. It is possible an absence of glycolysis limits the cycling rate of this pump, which would explain the shift in electrophysiological properties of ASER in the mutant. It is also possible that a glycolytically-derived metabolite could serve as a limiting factor in ASER function. For example, ASE neurons depend upon a conversion cycle between GTP and cGMP^45^, and loss of glycolysis could limit GTP availability and disrupt cGMP signaling. While future experiments will be required to establish the precise mechanisms by which glycolysis regulates ASER physiology, our data indicate that glycolysis is required to sustain membrane conductance and shape neuron-specific electrophysiological properties.

Our findings indicate that metabolism in neurons serves as both a hardwired determinant of functional specialization and a flexible substrate for plasticity. The divergent metabolic programs of ASEL and ASER arise in part from their cell-intrinsic transcriptional identities, but can also shift within minutes in response to neuronal activity, transient hypoxia or other physiological cues. This dual nature of metabolism as both a fixed trait and a tunable module is reflected in the distinct glycolytic baselines of the ASE neurons, as well as in the fast metabolic responses we observe during sensory stimulation. Because ASER responsiveness is known to be modulated by feeding state^46^, metabolic tuning provides a plausible node through which internal state may recalibrate sensory gain. Our findings support a model in which glycolysis not only dynamically adapts to changing metabolic demands as previously appreciated^9,16,37–39^, but also may function to shape neuronal responsiveness, tuning ionic conductance and modulating information flow within circuits.

A metabolic dimension of neuronal identity expands our understanding of neural circuit organization and function. By demonstrating that metabolic pathways actively shape neuronal physiology *in vivo*, this study reframes metabolism as a determinant—rather than a mere consequence—of neuronal signaling. Because resources are spatially distributed across cells and circuits, metabolism may impose constraints on the connectome that shapes how information is processed. It may be that metabolic “landscapes” modulate information flow in ways analogous to how synaptic weights and positions affect circuit dynamics. Incorporating metabolism into our framework of neuronal diversity thus provides new insights into how neurons perform computations and how these computations may change when metabolic pathways are perturbed.

## Acknowledgements

We thank the Hammarlund lab (Yale University, New Haven CT, USA) for sharing the codon-optimized construct of R-iLACCO1.2^26^. We thank the Yang lab (East China University of Science and Technology, Shanghai, CN) for sharing the sequence of SoNar^27^. We thank the Bringmann lab (TU-Dresden, Dresden, DE) for sharing the UNC-58 and TWK-18 constructs^43^. We thank the Albrecht lab (Worcester Polytechnic Institute, Worcester MA, USA) for the gift of microfluidics chambers for the salt stimulation experiments. We thank Gail Mandel, Lulu Cambronne, Miriam Goodman, Leonard Kaczmarek, Tony Hyman, the members of the Colón-Ramos lab for their thoughtful comments and discussions related to this project. Some strains were provided by the CGC, which is funded by NIH Office of Research Infrastructure Programs (P40 OD010440). This work was supported by National Institutes of Health grants to D.C.-R. (R35NS132156 and R01NS076558), to A.D.W. (K99AG083129), to A.J.M.W (R35GM122502 and R01DK068429), and to Z.-W.W (R01MH085927).

## Methods

### Molecular biology and constructs

Vectors for ASE expression of HYlight^25^, R-iLACCO1.2^26^, SoNar^27^, and JGCaMP8m^40^ were generated by inserting codon-optimized gene blocks of each sensor after a -3000 bp *flp-6* promoter. R-iLACCO1.2 and JGCaMP8m was placed in frame with a subsequent T2A peptide and TagBFP2 construct to enable ratiometric measurements over BFP fluorescence; SoNar was placed in frame with a T2A-mScarlet-I3 construct so that the 405 excitation can be used ratiometrically over mScarlet fluorescence. To label ASER, we used either mCherry (alongside HYlight and JGCaMP8m), mTagBFP (alongside SoNar), or mStayGold (alongside R-iLACCO1.2) driven by a -308 bp *gcy-5* promoter. Constructs for ectopic expression of hyperactive ion channels were made by inserting either UNC-58(L428F) or TWK-18(G165D) between the *flp-6* promoter and a T2A-mCherry.

### C. elegans maintenance and genetics

All worm strains were raised on nematode growth media at 20°C using the *Escherichia coli* strain OP50 as the sole food source on NGM/agar plates^47^, which have approximately 51 mM of Na^+^ and 53 mM of Cl^-^. Analyses were performed on day-old adults by picking animals at the L4 larval stage to seeded NGM plates and performing experiments within 16-20 hours. The *C. elegans* Bristol strain N2 was used as wild-type controls. A list of strains used in this study can be found in table S3.

### Transgenics

Transgenic strains were made by standard germline injection techniques; injection mixes of plasmids were normalized to 100 ng/uL of total DNA concentration using 100 kb ladder (New England Biolabs, Inc) as DNA filler. The strain expressing ASE::HYlight, DCR9228 (*olaEx5471*), was integrated using UV integration and outcrossed three generations to generate DCR9288 (*olaIs141*) and DCR9299 (*olaIs142*). CRISPR-Cas9 was used to generate genetic knockouts for *ldh-1(ola514)* based on published protocols^48^. Cut sites adjacent to the 5’ and 3’ sites of the gene were ordered from Horizon Discovery as modified crRNA oligos (5’-GATATCTATTAAAAATGACA-3’ and 5’-GTTCGATGACTGAAGGAAAA-3’ respectively). Worms from the F1 generation were singled and sequenced for the *ldh-1* deletion and the resulting homozygous progeny were outcrossed three times prior to use.

### Microfluidics and microscopy

Imaging was done by placing an 8% imaging pad of agarose in water on top of a PDMS microfluidics device designed for controlling local atmospheric conditions of worms^34^. Worms were placed within a 3 µL drop of imaging buffer (5 mM levamisole, 25 mM potassium phosphate pH 6, 1 mM calcium chloride, 1 mM magnesium sulfate, 50 mM NaCl, 102 mM glycerol). This buffer contained glycerol to increase the osmolality of the solution to approximately 260 mOsm to hold osmolality constant across NaCl concentrations ranging from 0 mM to 100 mM as required. 15-25 worms were placed in this drop without washing and allowed to sit for 10 minutes, covered with a plastic petri dish lid to prevent evaporation, before applying a No. 1.5 coverslip. This allowed the levamisole to take full effect and led to worms preferentially being on their backs when a coverslip is applied, allowing simultaneous visualization of both ASER and ASEL. For the identity swap experiments, the right and left neurons were identified based on their position in the left or right side of the animal, which was determined relative to the dorsal/ventral and anterior/posterior axes.

All imaging was performed on a Nikon Ti2 + CSU-W1 spinning disk confocal microscope and captured with a Hamamatsu Orca-Fusion BT CMOS camera at 16-bit pixel depth. After mounting, flow of compressed air was initiated immediately to prevent unwanted hypoxic conditions under the coverslip, and a holding period of 5 minutes was used to allow for thermal equilibration. During imaging, a manual valve was used to switch between air and pure compressed nitrogen gas at the same flow rate to induce hypoxic conditions.

Optimization of imaging parameters was done as previously described^15^. Conditions for each sensor were selected based on negative control experiments as available (e.g., HYlight-RA, or the *ldh-1* mutant for R-iLACCO1.2) by adjusting laser power settings to generate a mean ratio of approximately 1.0-1.5x for the appropriate wavelengths, using approximately 50-200 ms exposure times to exceed an independent signal:noise ratio for each channel of approximately 20-25. Images were captured for experiments between 0.2-1 Hz.

For determining parameters of the glycolysis stress test, values for basal glycolysis were defined as the HYlight signal before hypoxia is applied. Each experiment consisted of animals mounted onto a microfluidics device, exposed to a 1 min period with normal air, followed by a 5 minute period under pure nitrogen gas. We first used animals expressing the negative control HYlight-RA construct, and calculated the baseline signal averaged over the last 30 seconds under hypoxia (4.5-5 minutes). We then repeated this experiment with animals expressing HYlight and averaged the HYlight response over the last 30 seconds of the experiment. Glycolytic capacity was calculated as the difference in the response for each neuron in the individual animals examined relative to the mean value of HYlight-RA in either ASER or ASEL. Glycolytic reserve was defined as the difference between this same 30 second window in hypoxia relative to the first 30 seconds of the experiment during normoxia calculated for each individual neuron.

### Salt stimulation experiments

Experiments using salt flow microfluidics were performed by modifying the system and methods previously described^16^. Two NE-1002X syringe pumps (New Era Pump Systems, Inc., Farmingdale, NY) were loaded with variants of standard imaging buffer: a no-salt variant (NGM-0: 5 mM levamisole, 25 mM potassium phosphate pH 6, 1 mM calcium chloride, 1 mM magnesium sulfate, 202 mM glycerol) and a high-salt variant (NGM-100: 5 mM levamisole, 25 mM potassium phosphate pH 6, 1 mM calcium chloride, 1 mM magnesium sulfate, 100 mM NaCl, 2 mM glycerol). These were connected to a 600 µm passive microfluidics herringbone mixer (Darwin Microfluidics, Inc., Paris France; #CS-10001930) and flow-through conductivity electrode (Microelectrodes, Inc., Bedford NH; #16-900) in series. Tubing lengths were minimized to limit dead volume upon pump switching. This was connected to the input of a P10 PDMS microfluidics chamber^49^ and subsequent waste line. Control of the pump system was via a custom Qt6/Python interface^50^ written for this purpose; an improved version^51^ was subsequently built in C++, with GPT-4o (OpenAI, April 2025 version) and Claude Sonnet 4.5 (Anthropic, Sep 2025 version) used to assist with the code rewrite. Measurement of conductivity was achieved by continuous monitoring with a Orion Star A210 (Thermo Scientific, Inc) by using this custom interface software. The software is available on Github at https://github.com/adwolfe/PumpController.

Experiments for stimulating the ASE neurons were done by first anaesthetizing worms in imaging buffer containing either 0 mM NaCl or 50 mM NaCl, depending on chosen starting condition. Worms were washed in a 100 uL drop of imaging buffer on an unseeded NGM plate, then moved to a second 100 uL drop of imaging buffer for incubation for 10 minutes. A 1 mL syringe filled with the same imaging buffer was then used to aspirate the worms and inject them into the PDMS device worm chamber while flowing the starting buffer condition via the pumps. For comparing the change in glycolysis that occurred upon neuronal activation (Fig. 3B), a paradigm was selected that elicited a comparable calcium response in both cells. Worms were held at 0 mM NaCl, and then a 2 minute pulse of 50 mM NaCl was given, followed by a return to 0 mM NaCl. ASEL and ASER responded upon the increase and decrease of salt concentrations, respectively. For observing the effects of *pfk-1.1(ola458),* the paradigm was selected to maximize the change in calcium upon activation for either neuron. For ASEL, the same paradigm was used as before; for ASER, worms were held in 50 mM NaCl buffer, and then exposed to a 75 mM NaCl pulse for 2 minutes followed by a drop to 25 mM NaCl for 2 minutes.

### Metabolic network modeling

Single-cell RNA-seq data from the CeNGEN project^31^ were aggregated into pseudo-bulk profiles using previously-described methods^52^, resulting in mRNA expression data in transcripts per million (TPM). All cell types represented by fewer than 100 single cells were excluded. For comparison, non-neuronal cell types with more than 500 sampled cells were included, with the exception of intestine, due to its unique metabolic role and direct access to degraded bacterial biomass^53^, and body-wall muscle (anterior), which is physiologically similar to body-wall muscle and therefore redundant. The final dataset comprised 103 neuronal cell types, including 26 sensory neurons, and 10 non-neuronal cell types. These data were integrated with the *C. elegans* metabolic network model iCEL1314 through the MERGE computational pipeline^32^ as explained below.

Network-level optimized flux distributions were generated for each cell type using the iMAT++ algorithm with the following settings: (i) first, gene expression data were discretized into expression states using CatExp^32^. (ii) The iCEL1314 model was used with its default compartment structure. (iii) To avoid enforcing full biomass production in specialized cell types that synthesize only a subset of biomass components, the biomass assembly pathway was relaxed by introducing drain reactions consuming major biomass precursors (e.g., collagen and glycans). (iv) The model was constrained to allow uptake of nutrients that can be supplied by the intestine or are required for specific reactions to carry flux. Uptake of metabolites of non-bacterial origin and exchange of intermediate metabolites (*e.g*., TCA-cycle intermediates such as isocitrate) were discouraged by assigning the corresponding exchange reactions to the lowly expressed reaction set. Metabolites that are not part of the endogenous *C. elegans* metabolome^54^ and may occur only as supplements, such as testosterone, were blocked using hard constraints. (v) Flux variability analysis (FVA) was performed where needed to determine the minimum and maximum feasible flux for each reaction, thereby characterizing the alternative solution space not captured by the optimized flux values.

Flux potential analysis was performed using the updated eFPA algorithm^30^ with the following settings: (i) first, the modified iCEL1314 model used for iMAT++ analysis was retained. (ii) A default distance boundary of six reactions was used to define the local network neighborhood prioritized for expression integration^30^. (iii) To discourage uptake and exchange of uncommon or intermediate metabolites (see above), the special penalties option of eFPA was used, penalizing the relevant boundary reactions by a factor of 25. (iv) For each cell type, reactions identified by FVA as incapable of carrying flux in one or both directions were constrained accordingly by adjusting reaction bounds, thereby preventing eFPA from using flux values outside the iMAT++ solution space.

### Electrophysiology

Electrophysiological recordings were performed on Day 1 adult *C. elegans* hermaphrodites, using the strains DCR9288 for wild type and DCR9887 for *pfk-1.1(ola458)* experiments. For each experiment, an animal was immobilized by applying a small drop of 3M VetbondTM (1469SB) to the dorsal head region on a Sylgard-coated coverslip. A diamond dissecting tool (72028, Electron Microscopy Sciences, Hatfield, PA, USA) was used to make an incision through the glued cuticle. The resulting cuticle flap was pulled open and secured to the coverslip using GLUture (Zoetis Inc., Kalamazoo, MI, USA). The pharynx was then gently displaced to expose ASER and ASEL sensory neurons. ASE neurons were identified based on green HYlight fluorescence (*flp-6* promoter) and their anatomical positions relative to the dorsal/ventral and anterior/posterior axes, and this identification was confirmed by the presence of ASER-specific mCherry fluorescence (*gcy-5* promoter).

Whole-cell current-clamp recordings were obtained from both ASER and ASEL in each animal. A series of current injection steps (-1.0 pA to +6.0 pA, 0.5 pA increments, 600 ms duration) were applied to evoke changes in membrane potential. To mitigate potential time-dependent effects, the sequence of ASER and ASEL recordings was alternated across animals. ASER was recorded first in three of five wild-type animals and three of six *pfk-1.1(ola458)* mutant animals. Data from animals in which only one of the two neurons was successfully recorded were excluded from analysis.

Recordings were performed using borosilicate glass electrodes with tip resistance of approximately 20 MΩ. Signals were filtered at 2 kHz and sampled at 10 kHz using a dPatch Digital Patch Clamp Amplifier controlled by SutterPatch software (Sutter Instruments). The extracellular (bath) solution contained (in mM) 50 NaCl, 90 N-methyl-d-glucamine (NMDG), 5 KCl, 5 CaCl₂, 1 MgCl₂, 11 dextrose, and 5 HEPES (pH adjusted to 7.3 with HCl). The intracellular (pipette) solution contained (in mM) 107 potassium gluconate (K-Glu), 15.5 KOH, 13 KCl, 0.25 calcium gluconate (Ca-Glu), 4 MgCl₂, 5 Tris, 36 sucrose, 5 EGTA, and 4 Na₂ATP (pH adjusted to 7.2 with 8 mM HCl).

Electrophysiological recordings were exported as ABF files for subsequent quantification of the RMP, as well as the rise time and 90% decay time of membrane potential responses to current injection, using ClampFit (Molecular Devices). The RMP was calculated as the average membrane potential during the initial 100-ms pre-pulse interval across all 15 current injection steps. Rise time was defined as the interval between the onset of the current injection and the time to peak membrane potential. The 90% decay time was measured from the termination of the current injection to the point at which the membrane potential had decayed to 10% of its amplitude. Input resistance was determined from the slope of a linear fit to the relationship between injected current and the steady-state membrane potential. Current-voltage relationships were constructed from the steady-state membrane voltage changes (the average membrane voltage over the last 500 ms of the 600-ms current pulse) elicited by the first five current injection steps (ranging from –1.0 pA to +1.0 pA in 0.5-pA increments).

---

**Figure S1:**
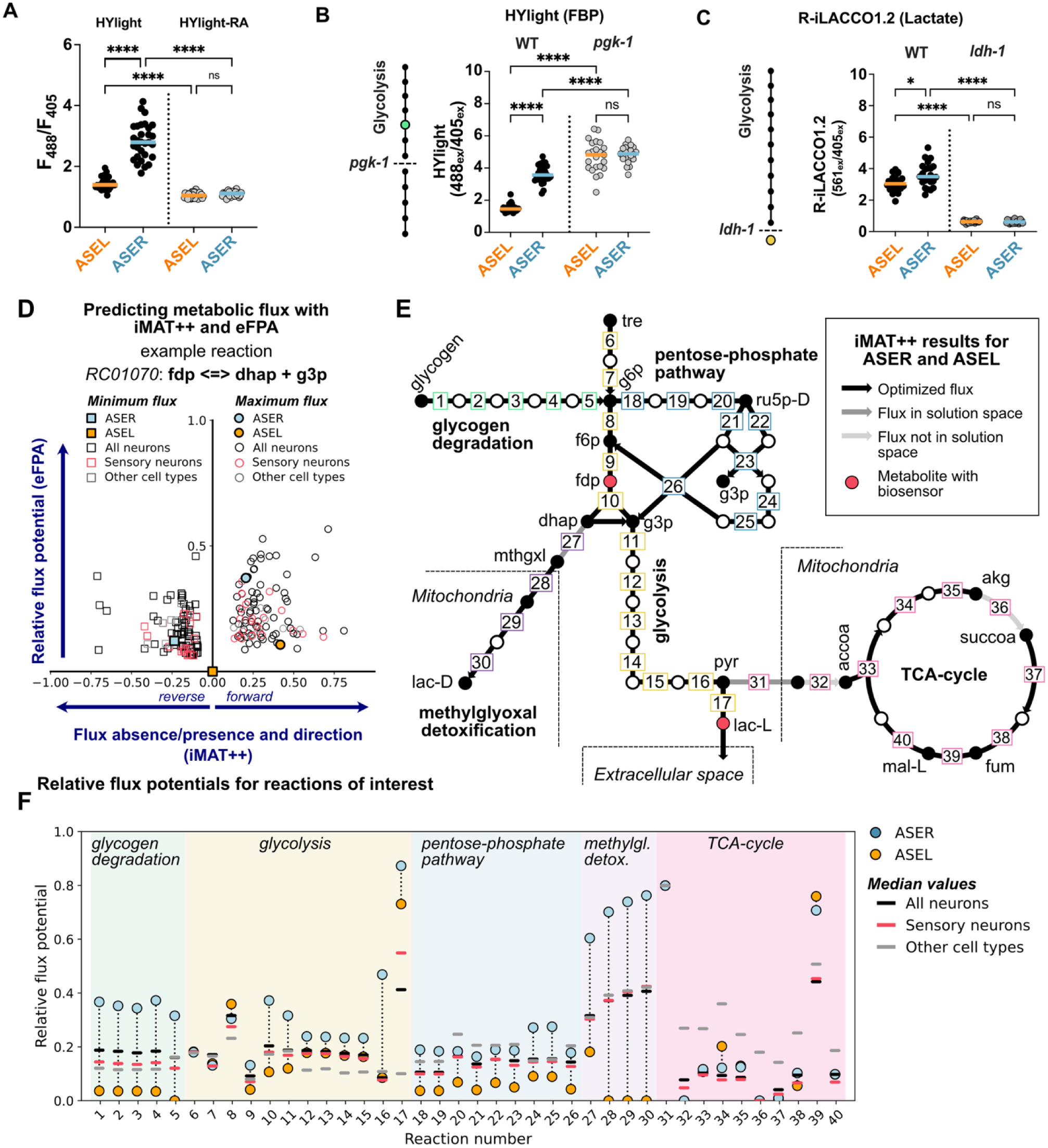
Supporting data for metabolic differences between ASER and ASEL. (A) Quantification of mean soma values comparing ASER and ASEL for HYlight (data taken from Fig. 1C) and the negative control variant, HYlight-RA. (B) HYlight measurements for FBP in wild-type vs *pgk-1(tm5613)* loss-of-function mutant animals. (C) R-iLACCO1.2 measurements for lactate in wild-type or *ldh-1(ola514)* knockout animals. (A-C) \**P*<0.5; \*\*\*\**P*<0.0001; ns, non-significant. Statistical details available in table S2. (D) Joint visualization of iMAT++ flux range (x-axis) and eFPA flux potential (y-axis) for the fructose-1,6-bisphosphate aldolase reaction across cell types. Each cell type is represented by two points corresponding to the minimum and maximum feasible flux values derived from flux variability analysis (FVA) of the iMAT++ solution space, together defining the allowable flux range. A positive maximum flux indicates support for the forward reaction direction, whereas a negative minimum flux indicates support for the reverse direction. ASER and ASEL are highlighted with filled markers. Values less than −1 and greater than 1 on the x-axis are omitted for clarity (<10% of data). (E) Reactions in pathways of interest with flux solutions inferred by iMAT++. Line color indicates whether fluxes in the indicated pathway direction (arrow orientation) are present in the optimized iMAT++ flux distribution (default solution) or fall within the allowable flux ranges (solution space) of ASER or ASEL based on iMAT++ predictions. FBP is denoted as fdp, consistent with the iCEL1314 metabolic network model. Reaction numbering corresponds to table S1 and panel (F). See table S1 for details. (F) Relative flux potentials from eFPA for the reactions shown in (E), with ASER and ASEL highlighted. Median values are shown for each cell-type group. Note that ASER is predicted to have an exceptionally high flux potential for glycolytic-adjacent pathways, including the methylglyoxal detoxification pathway, which detoxifies methylglyoxal generated primarily from triose phosphates under high glycolytic flux^55^.

**Figure S2:**
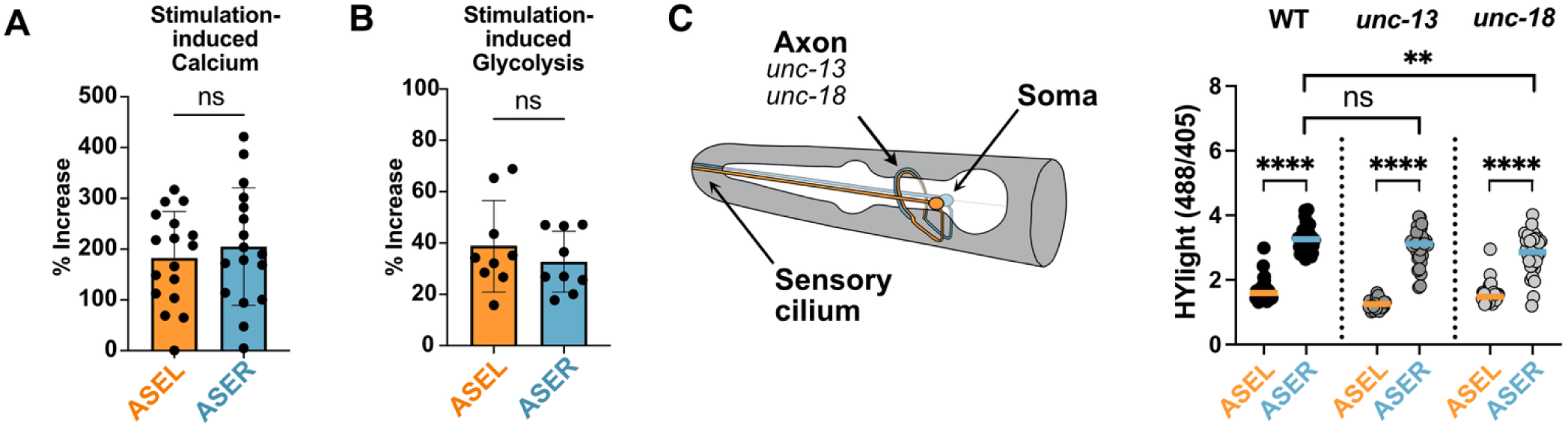
Supporting data for determining the source of metabolic differences in ASE neurons. (A) Comparison of the percent change for data shown in Fig. 3B-C. Left, the calculated percent change for the JGCaMP8m/BFP ratio (corresponding to Fig. 3B) from the mean peak value (16 second window centered on the peak maximum) for each animal relative to the 16 second window immediately preceding the response. (B) The same time points were used to determine the glycolysis response as shown by HYlight (Fig. 3C). (C) The genes *unc-13* and *unc-18* primarily function in synaptic vesicle release in the axon; right, the effect of *unc-13(s69)* and *unc-18(e234)* loss-of-function alleles on HYlight baselines of ASE neurons. \*\**P*<0.01; \*\*\*\**P*<0.0001; ns, non-significant. Statistical details available in table Ss2.

**Figure S3:**
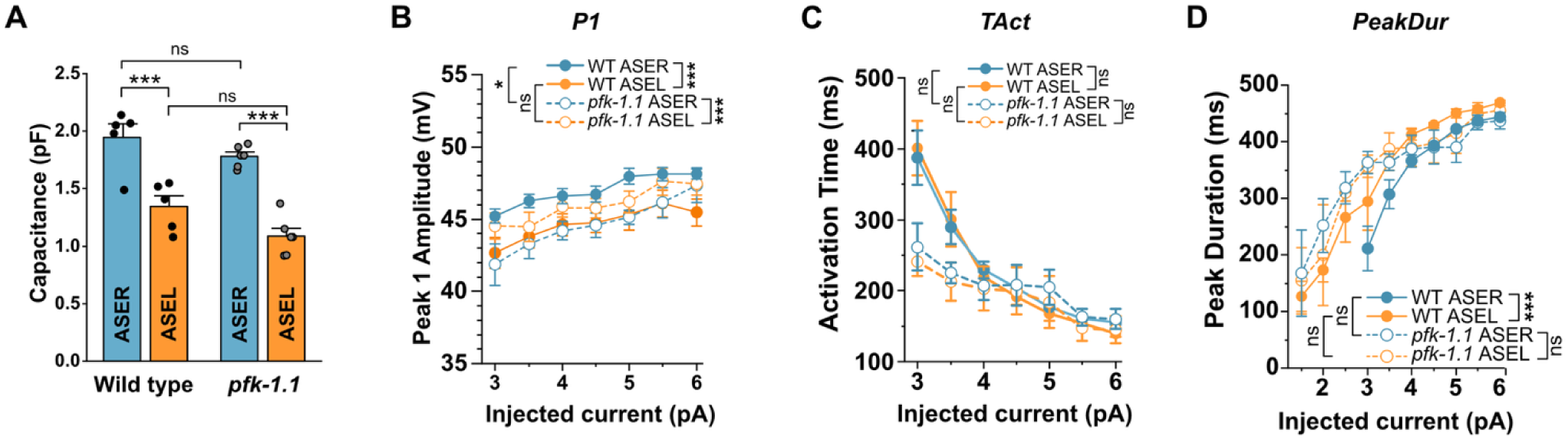
Supplementary electrophysiology of ASE neurons. (A) Comparisons of membrane capacitance between ASER and ASEL across wild-type and *pfk-1.1(ola458)* animals. (B-D) Comparisons of Peak 1 amplitude (B), activation time (C), and peak duration (D) of current-induced membrane voltage changes across neurons and genotypes. See Fig. 4E for definition of P1, Tact, and PeakDur. Data are shown as mean ± SEM. \**P*<0.5; \*\**P*<0.01; \*\*\**P*<0.001; ns, non-significant. Statistical details available in table S2.

**Table S1: Results of flux modeling for key metabolic pathways (see** Fig. 1H**, S1C-E)**

**Table S2: List of statistical methods and results.**

**Table S3: List of strains used in this study.**

## Notes

### Competing Interest Statement

The authors have declared no competing interest.

## References

1. Llinás, R. R. The intrinsic electrophysiological properties of mammalian neurons: insights into central nervous system function. Science 242, 1654–1664 (1988).

2. Marder, E. & Goaillard, J.-M. Variability, compensation and homeostasis in neuron and network function. Nat Rev Neurosci 7, 563–574 (2006).

3. Zeng, H. & Sanes, J. R. Neuronal cell-type classification: challenges, opportunities and the path forward. Nat Rev Neurosci 18, 530–546 (2017).

4. Attwell, D. & Laughlin, S. B. An Energy Budget for Signaling in the Grey Matter of the Brain. Journal of Cerebral Blood Flow & Metabolism 21, 1133–1145 (2001).

5. Harris, J. J., Jolivet, R. & Attwell, D. Synaptic Energy Use and Supply. Neuron 75, 762– 777 (2012).

6. Li, S. & Sheng, Z.-H. Energy matters: presynaptic metabolism and the maintenance of synaptic transmission. Nat Rev Neurosci 23, 4–22 (2022).

7. Qin, C. et al. Signaling pathways involved in ischemic stroke: molecular mechanisms and therapeutic interventions. Sig Transduct Target Ther 7, 215 (2022).

8. Ni, A. & Ernst, C. Evidence That Substantia Nigra Pars Compacta Dopaminergic Neurons Are Selectively Vulnerable to Oxidative Stress Because They Are Highly Metabolically Active. Front Cell Neurosci 16, 826193 (2022).

9. Ashrafi, G., Wu, Z., Farrell, R. J. & Ryan, T. A. GLUT4 Mobilization Supports Energetic Demands of Active Synapses. Neuron 93, 606–615 3 (2017).

10. Yellen, G. Fueling thought: Management of glycolysis and oxidative phosphorylation in neuronal metabolism. Journal of Cell Biology 217, 2235–2246 (2018).

11. Dienel, G. A. Brain Glucose Metabolism: Integration of Energetics with Function. Physiological Reviews 99, 949–1045 (2019).

12. Mills, E. L. et al. Succinate Dehydrogenase Supports Metabolic Repurposing of Mitochondria to Drive Inflammatory Macrophages. Cell 167, 457–470.e13 (2016).

13. Bourdeau Julien, I., Sephton, C. F. & Dutchak, P. A. Metabolic Networks Influencing Skeletal Muscle Fiber Composition. Front Cell Dev Biol 6, 125 (2018).

14. Wong-Riley, M. Energy metabolism of the visual system. Eye Brain 2, 99–116 (2010).

15. Wolfe, A. D. et al. Local and dynamic regulation of neuronal glycolysis in vivo. Proceedings of the National Academy of Sciences 121, e2314699121 (2024).

16. Singh, M. et al. Glycogen supports glycolytic plasticity in neurons. Proceedings of the National Academy of Sciences 122, e2509003122 (2025).

17. Hobert, O., Johnston, R. J. & Chang, S. Left–right asymmetry in the nervous system: the Caenorhabditis elegans model. Nat Rev Neurosci 3, 629–640 (2002).

18. Chang, S., Johnston, R. J. & Hobert, O. A transcriptional regulatory cascade that controls left/right asymmetry in chemosensory neurons of C. elegans. Genes Dev. 17, 2123– 2137 (2003).

19. Johnston, R. J. & Hobert, O. A microRNA controlling left/right neuronal asymmetry in Caenorhabditis elegans. Nature 426, 845–849 (2003).

20. Pierce-Shimomura, J. T., Faumont, S., Gaston, M. R., Pearson, B. J. & Lockery, S. R. The homeobox gene lim-6 is required for distinct chemosensory representations in C. elegans. Nature 410, 694–698 (2001).

21. Suzuki, H. et al. Functional asymmetry in Caenorhabditis elegans taste neurons and its computational role in chemotaxis. Nature 454, 114–117 (2008).

22. Hiroki, S. et al. Molecular encoding and synaptic decoding of context during salt chemotaxis in C. elegans. Nat Commun 13, 2928 (2022).

23. Ortiz, C. O. et al. Lateralized Gustatory Behavior of C. elegans Is Controlled by Specific Receptor-Type Guanylyl Cyclases. Current Biology 19, 996–1004 (2009).

24. Luo, L. et al. Dynamic Encoding of Perception, Memory, and Movement in a C. elegans Chemotaxis Circuit. Neuron 82, 1115–1128 (2014).

25. Koberstein, J. N. et al. Monitoring glycolytic dynamics in single cells using a fluorescent biosensor for fructose 1,6-bisphosphate. Proceedings of the National Academy of Sciences 119, e2204407119 (2022).

26. Nasu, Y. et al. Lactate biosensors for spectrally and spatially multiplexed fluorescence imaging. Nat Commun 14, 6598 (2023).

27. Zhao, Y. et al. SoNar, a Highly Responsive NAD+/NADH Sensor, Allows High-Throughput Metabolic Screening of Anti-tumor Agents. Cell Metabolism 21, 777–789 (2015).

28. Zhang, H. et al. A systems-level, semi-quantitative landscape of metabolic flux in C. elegans. Nature 640, 194–202 (2025).

29. Li, X. et al. Systems-level design principles of metabolic rewiring in an animal. Nature 640, 203–211 (2025).

30. Li, X., Walhout, A. J. M. & Yilmaz, L. S. Enhanced flux potential analysis links changes in enzyme expression to metabolic flux. Mol Syst Biol 21, 413–445 (2025).

31. Taylor, S. R. et al. Molecular topography of an entire nervous system. Cell 184, 4329–4347.e23 (2021).

32. Yilmaz, L. S. et al. Modeling tissue-relevant Caenorhabditis elegans metabolism at network, pathway, reaction, and metabolite levels. Mol Syst Biol 16, MSB209649 (2020).

33. Yoo, I., Ahn, I., Lee, J. & Lee, N. Extracellular flux assay (Seahorse assay): Diverse applications in metabolic research across biological disciplines. Molecules and Cells 47, 100095 (2024).

34. Jang, S. et al. Phosphofructokinase relocalizes into subcellular compartments with liquid-like properties in vivo. Biophysical Journal 120, 1170–1186 (2021).

35. Poole, R. J., Bashllari, E., Cochella, L., Flowers, E. B. & Hobert, O. A Genome-Wide RNAi Screen for Factors Involved in Neuronal Specification in Caenorhabditis elegans. PLOS Genetics 7, e1002109 (2011).

36. Sarin, S. et al. Genetic Screens for Caenorhabditis elegans Mutants Defective in Left/Right Asymmetric Neuronal Fate Specification. Genetics 176, 2109–2130 (2007).

37. Rangaraju, V., Calloway, N. & Ryan, T. A. Activity-Driven Local ATP Synthesis Is Required for Synaptic Function. Cell 156, 825–835 (2014).

38. Díaz-García, C. M. et al. Neuronal Stimulation Triggers Neuronal Glycolysis and Not Lactate Uptake. Cell Metabolism 26, 361–374.e4 (2017).

39. Díaz-García, C. M. et al. The distinct roles of calcium in rapid control of neuronal glycolysis and the tricarboxylic acid cycle. eLife 10, e64821 (2021).

40. Zhang, Y. et al. Fast and sensitive GCaMP calcium indicators for imaging neural populations. Nature 615, 884–891 (2023).

41. Pulido, C. & Ryan, T. A. Synaptic vesicle pools are a major hidden resting metabolic burden of nerve terminals. Science Advances 7, eabi9027 (2021).

42. Andrini, O. et al. Constitutive sodium permeability in a Caenorhabditis elegans two-pore domain potassium channel. Proceedings of the National Academy of Sciences 121, e2400650121 (2024).

43. Busack, I. & Bringmann, H. A sleep-active neuron can promote survival while sleep behavior is disturbed. PLOS Genetics 19, e1010665 (2023).

44. Meyer, D. J., Díaz-García, C. M., Nathwani, N., Rahman, M. & Yellen, G. The Na+/K+ pump dominates control of glycolysis in hippocampal dentate granule cells. eLife 11, 81645 (2022).

45. Smith, H. K. et al. Defining Specificity Determinants of cGMP Mediated Gustatory Sensory Transduction in Caenorhabditis elegans. Genetics 194, 885–901 (2013).

46. Tomioka, M. et al. The Insulin/PI 3-Kinase Pathway Regulates Salt Chemotaxis Learning in *Caenorhabditis elegans*. Neuron 51, 613–625 (2006).

47. Brenner, S. THE GENETICS OF CAENORHABDITIS ELEGANS. Genetics 77, 71–94 (1974).

48. Dickinson, D. J. & Goldstein, B. CRISPR-Based Methods for Caenorhabditis elegans Genome Engineering. Genetics 202, 885–901 (2016).

49. Albrecht, D. R. & Bargmann, C. I. High-content behavioral analysis of Caenorhabditis elegans in precise spatiotemporal chemical environments. Nat Methods 8, 599–605 (2011).

50. Wolfe, A. D. PyPumpController: v1.6.1. Zenodo 10.5281/zenodo.18669953 (2026).

51. Wolfe, A. PumpController: v1.2.1b. Zenodo 10.5281/zenodo.18670672 (2026).

52. Cao, J. et al. Comprehensive single-cell transcriptional profiling of a multicellular organism. Science 357, 661–667 (2017).

53. McGhee, J. D. The C. elegans intestine. in WormBook: The Online Review of C. elegans Biology [Internet] (WormBook, 2007).

54. WormBase 2024: status and transitioning to Alliance infrastructure - PubMed. https://pubmed.ncbi.nlm.nih.gov/38573366/.

55. Allaman, I., Bélanger, M. & Magistretti, P. J. Methylglyoxal, the dark side of glycolysis. Front Neurosci 9, 23 (2015).

